# Molecular mechanisms associated with clustered lesion-induced impairment of 8-oxoG recognition by the human glycosylase OGG1

**DOI:** 10.1101/2021.09.23.461474

**Authors:** Tao Jiang, Antonio Monari, Elise Dumont, Emmanuelle Bignon

## Abstract

The 8-oxo-7,8-dihydroguanine, referred to as 8-oxoG, is a highly mutagenic DNA lesion that can provoke the appearance of mismatches if it escapes the DNA Damage Response. The specific recognition of its structural signature by the hOGG1 glycosylase is the first step along the Base Excision Repair pathway, that ensures the integrity of the genome by preventing the emergence of mutations. 8-oxoG formation, structural features and repair have been the matter of extensive research and more recently this active field of research expended to the more complicated case of 8-oxoG within clustered lesions. Indeed, the presence of a second lesion within 1 or 2 helix turns can dramatically impact the repair yields of 8-oxoG by glycosylases. In this work, we use *μ*s-range molecular dynamics simulations and machine learning-based post-analysis to explore the molecular mechanisms associated with the recognition of 8-oxoG by hOGG1 when embedded in a multiple lesions site with a mismatch in 5’ or 3’. We delineate the stiffening of the DNA-protein interactions upon the presence of the mismatches, and rationalize the much lower repair yields reported with a 5’ mismatch by describing the perturbation of 8-oxoG structural features upon addition of an adjacent lesion.

## 1. Introduction

8-oxo-7,8-dihydroguanine (8-oxoG) is the most common oxidatively-generated DNA damage and involves at least 1 over 10^6^ of the total pool of guanines in human cells [1]. It is mostly generated by oxidative stress, either upon hydration of the guanine radical cation or by reaction of hydroxyl radical (·OH) with guanine [2,3]. Additionally, it may also be generated by UV irradiation, most notably via the activation of singlet oxygen by photosensitizers. The high carcinogenic potential of 8-oxoG arises from its capacity to bypass DNA polymerases checkpoints, leading to the incorporation of an adenine in the nascent strand and ultimately to G:C → A:T mutations [4,5]. When 8-oxoG is not isolated but is, instead, embedded in multiple damaged sites, i.e. multiple spatially close DNA lesions which are particularly challenging for DNA damage response, its mutagenicity is even higher [6–9]. Of note, multiple damaged sites within one or two helical turns are referred to as clustered lesions [10].

To overcome the impact of 8-oxoG on DNA integrity, this lesion is efficiently repaired by Base Excision Repair enzymes [11,12]. Human OGG1 (hOGG1) is the first enzyme involved in 8-oxoG repair. It catalyzes the hydrolysis of the N-glycosidic bond and the subsequent *β*-elimination of the 3’ phosphate of the resulting abasic site. The molecular mechanisms driving 8-oxoG recognition by OGG1 constitute a complex matter of research [13], and a recent joint cryo-EM/*in silico* study of hOGG1 bound to an oligonucleotide harboring an intrahelical 8-oxoG provides crucial insights into key-residues involved in the 8-oxoG extrusion process [14]. A most important and conserved structural feature of OGG1 is the N149-N150-N151 motif, which plays a major role in interacting with the DNA helix and stabilizing the 8-oxoG-extruded structure. Especially, the N149, which interacts directly with the intra-helical 8-oxoG in the minor groove, fills the gap resulting from the absence of the oxidized nucleobase which is extruded through the major groove and stabilizes the double helix - see Figure 1-A. Two arginines (R154 and R204) interact with both N149 and the facing orphan cysteine to form a stable hydrogen-bond network. Besides, a tyrosine (Y203) partially inserts into the helix to maintain the helix bend by *π*-stacking with the nucleobase in the 3’ position of the orphan cysteine.

**Figure 1.**
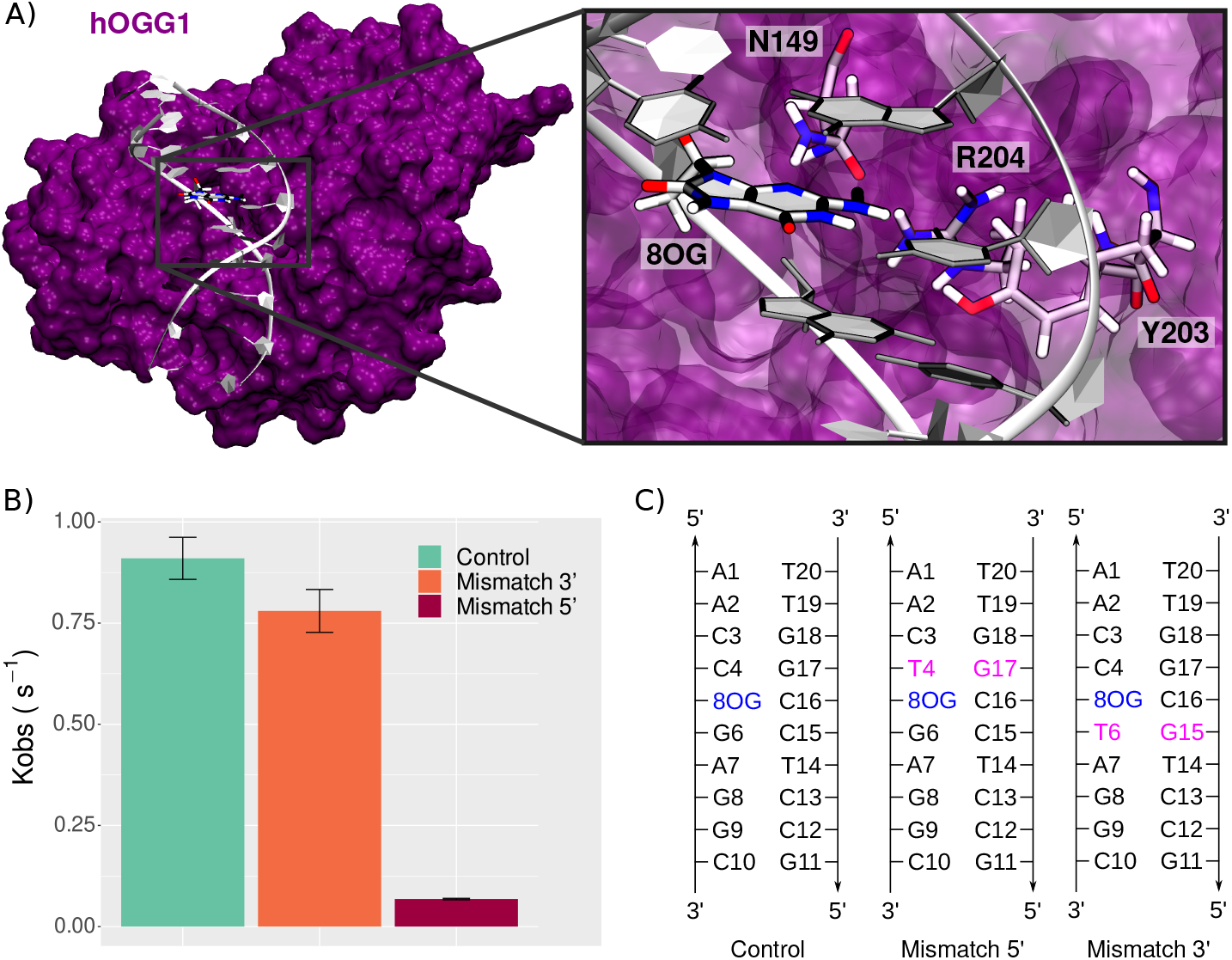
8oxoG processing by hOGG1. (**A**) Structure of the hOGG1 in interaction with the 10-bp DNA duplex bearing 8oxoG at position 5. The magnified section highlights the presence of three amino acids important for 8-oxoG extrusion: N149, Y203 and R204. For sake of clarity, backbone atoms are not depicted. (**B**) Repair rates of 8-oxoG by hOGG1 (control) reported by Sassa et al. upon presence of clustered lesions, i.e. a mismatch in 3’ or in 5’ of 8-oxoG [18]. (**C**) DNA sequences used in our simulations to probe the effect of a mismatch on the structural behavior of the hOGG1:DNA complex.

Upon the presence of clustered lesions hOGG1 repair rates can undergo dramatic decrease, depending on the nature and the position of the second lesion [15–17]. Sassa et al. investigated sequence effects and the presence of adjacent lesions on hOGG1 repair rates [18]. They highlighted the relative tolerance of hOGG1 concerning sequence effects, with only a slight decrease when 8-oxoG is flanked by either 5’ C, A, T or G. However, hOGG1 efficiency is dramatically impacted by the presence of a mismatch at 5’ position of 8-oxoG, with repair rates reduced by up to 100-fold [18] - see Figure 1-B. Interestingly, a similar mismatch placed 3’ of the lesion has a negligible effect on hOGG1 efficiency.

Owing to *μ*s molecular dynamics (MD) simulations, we investigated the effect of the presence of a mismatch adjacent to 8-oxoG on the hOGG1:DNA interaction, hence the damaged recognition and processing. Our results reveal contrasted fingerprints of the hOGG1:DNA interaction upon the presence of clustered mismatch and 8oxoG. The perturbation of the Watson-Crick network upon the presence of an additional mismatch leads to stronger interactions with the surrounding amino acids, which might disfavor the lesion extrusion. Machine Learning post-analysis highlight that hOGG1 region that are in contact with the surroundings of the lesion site, such as the highly-conserved NNN motif, stiffen upon the addition of a mismatch adjacent to 8-oxoG. Interestingly, the position of the mismatch in 5’ or 3’ affects only in a subtle way the helical properties, yet the thorough analysis of the DNA-protein interaction network reveals contrasted patterns that might rationalize the difference in repair rates. Thus, our simulations bring insights into the molecular mechanisms underlying the perturbation of hOGG1 efficiency upon the presence of clustered lesions, and especially help rationalizing the sequence effects highlighted by several previous experimental works.

## 2. Materials and Methods

### 2.1. MD simulations

All MD simulations were performed with NAMD2.14 [19], on systems set up using AmberTools18 [20]. The ff14SB force field and the bsc1 corrections for DNA backbone parameters were employed [21,22], and parameters for the 8oxoG non-canonical nucleobase were taken from previous works [23]. The systems were built from the experimental structure of OGG1 interacting with intrahelical 8oxoG-bearing 9bp DNA duplex PDB ID 6W13 [14]. To avoid edge effects, the DNA duplex was extended to 10bp. The DNA sequence was modified to match the ones used by Sassa et al. to allow the direct comparison of the MD results with the experiments [18] - see Figure 1-C. The missing loop 79-83 was reconstructed using the SwissModel server [24], the Y207C and C253W mutations added for crystallization purposes were reversed, and crystallographic waters were kept as such. Each system was soaked in an orthorhombic TIP3P water box with a 12.0 Å buffer and neutralized with K^+^cations, for a total of ~60,000 atoms that were minimized in three runs of 30,000 steps with decreasing restraints on backbone atoms. Each minimization run was coupled with a 4 ns equilibration run at 300K to ensure a smooth relaxation. A 400 ns production was then carried out to provide sampling for structural analysis. The temperature was maintained at 300 K using a Langevin Thermostat with a collision frequency *γ*–ln set to 1 ps^−1^. Electrostatics were handled using the Particle Mesh Ewald method [25] and a 9 Å cutoff was used for non-bonded interactions. The hydrogen-mass repartitioning (HMR) approach was used to allow stable simulations with a time steps of 4 fs, and simulations information were saved every 40 ps. Two replicates were performed for each system, resulting in a total of 2.4 *μ*s sampling.

### 2.2. Structural analysis

Structural parameters of the DNA duplex were computed using Curves+ [26] on 1000 frames extracted from each replicate. The other descriptors monitoring and the clustering were carried out using the cpptraj module of AmberTools18 [20]. The clustering of the structures were performed based on RMSD deviations of the 5-bp centered on the 8oxoG and the amino acids within 7 Å of the 8-oxoG. Plots, pictures and figures were generated using the ggplot2 package of R3.6.3 [27,28], VMD [29] and Inkscape (https://inkscape.org/).

The assessment of the protein flexibility was performed using a machine learning protocol proposed by Fleetwood and coworkers [30], in-house implemented and previously successfully used on the MutM bacterial analog of OGG1 [23].

## 3. Results

### 3.1. Protein-DNA interaction network

The interaction previously observed between hOGG1 and damaged oligonucleotides are correctly reproduced in our simulations. Hereafter, we describe the key residues interacting with the damaged region and forming the protein-DNA contact surface, and we scrutinize the perturbations of this network upon the presence of a 3’ or 5’ mismatch.

#### 3.1.1. Interactions in proximity of the damaged site

Several amino acids surrounding the damage site have been previously identified as being crucial for 8-oxoG extrusion: N149, R154, R204 and Y203 - see Figure 2.

**Figure 2.**
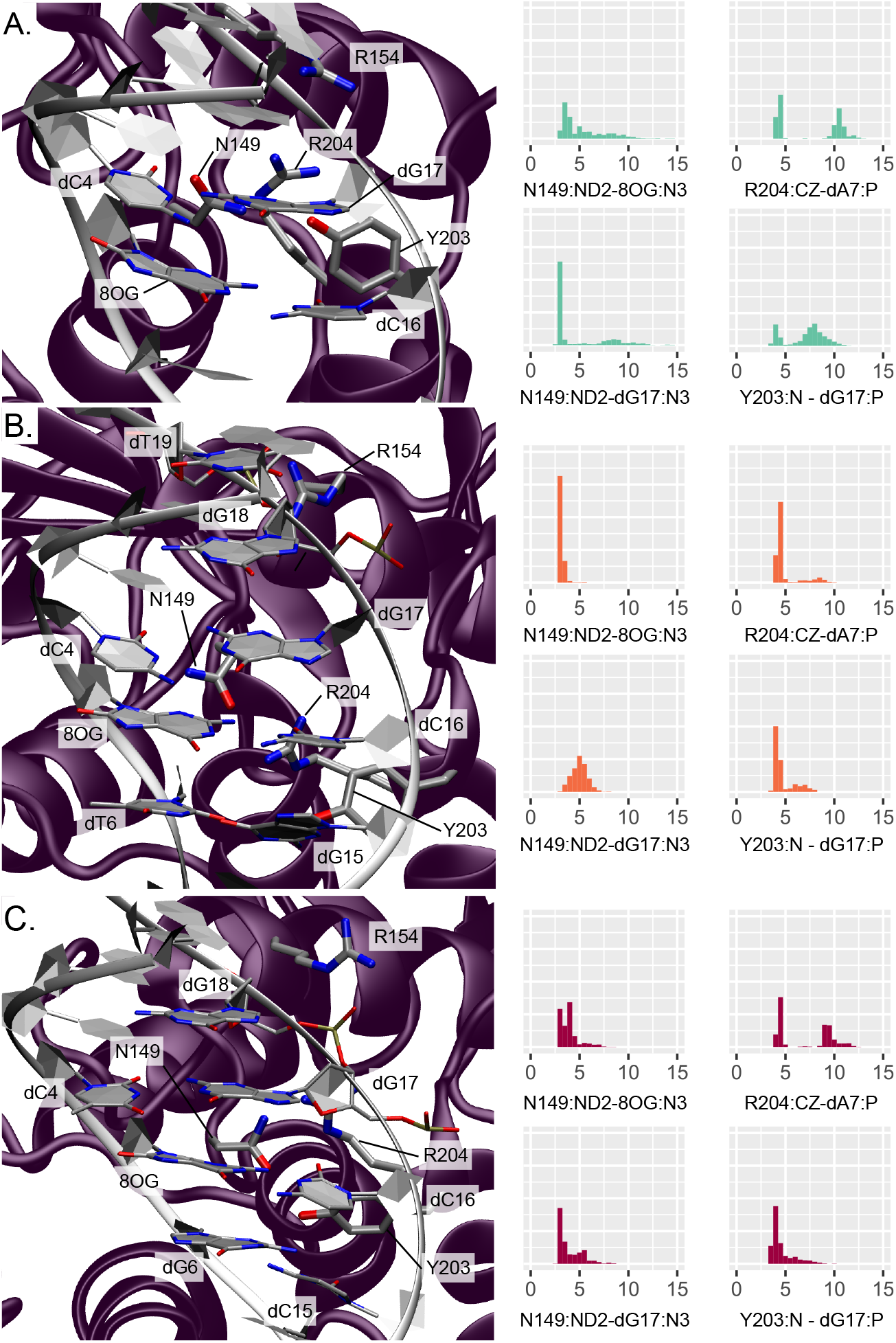
Representation and distribution of the key-interactions around the lesion site. **A.** for an isolated 8-oxoG, **B.** for clustered 8-oxoG and 3’ mismatch, **C.** for clustered 8-oxoG and 5’ mismatch. The protein is depicted in purple and the DNA helix in white, the interacting residues side chain are displayed in licorice with thicker tubes for amino acids. The histograms reflect the normalized distribution of the key-distances in Å.

The presence of the nearby N149 belonging to the NNN motif is especially important as this residue fills the gap in the helix left by the 8-oxoG extrusion, thus stabilizing the oligonucleotide. Upon the presence of an isolated 8-oxoG, our simulations show that this residue remains close to the damage site in the minor groove but interacts only transiently within the lesion surroundings, situating at distances from 8-oxoG and dG17 of 5.27 ± 2.20 Å (N149:ND2-8OG:N3) and 4.75 ± 2.85 Å (N149:ND2-dG17:N3), respectively. This is in agreement with the recent study by Shigdel et al. showing that N149 interacts with both nucleobases of the damaged base-pair [14].

The two arginines R154 and R204 also play an important role in stabilizing the DNA helix after 8-oxoG extrusion from the duplex, by interacting with the inserted N149 and the facing orphan base. Along our MD simulations, R154 makes contacts with dT19 and dG18 backbone atoms (R154:CZ-dT19:P and -dG18:P distances of 6.49 ± 2.33 Å and 5.42 ± 1.56 Å, respectively), keeping it close enough to the lesion site for further stabilization. R204 interacts with the target strand, either in the minor groove with the dG6 nucleobase (R204:CZ-dG6:N3 at 8.55 ± 2.48 Å) or with the dA7 phosphate group below (R204:CZ-dA7:P at 7.42 ± 3.16 Å). Finally, the stabilization of the 8-oxoG extruded structure also involves a partial insertion of Y203 in 3’ of the orphan base (dC16). In our simulations, Y203 stays nearby the minor groove, interacting mainly with dG17 on the facing strand through its backbone atom (Y203:N-dG17:P distance of 7.07 ± 1.93 Å) but also transiently with the target strand with either the dA7 sugar (Y203:HH-dA7:O4’ distance of 5.65 ± 2.48 Å) or the dG6 nucleobase (Y203:HH-dG6:N3 distance of 6.25 ± 2.47 Å).

Upon addition of an adjacent mismatch, the above-described interaction patterns exhibit perturbations that results in a stiffening of the DNA-protein contact region around the lesion, the position in 5’ or 3’ to the lesion dictating the new interaction network.

When the secondary mismatch is located in 3’, N149 interacts more tightly with 8-oxoG and dG17 (3.12 ± 0.28 Å and 4.98 ± 0.77 Å, respectively). Besides, R154 gets closer to dT19 phosphate (5.45 ± 1.46 Å) at the expense of dG18 (6.56 ± 1.84 Å) and R204 gets involved in more exclusive hydrogen bonding with dA7 phosphate (R204:CZ-dA7:P at 4.89 ± 1.32 Å). Similarly, Y203 also forms more stable interactions with the DNA duplex: it gets mainly involved in persistent hydrogen bonding with dG15 from the mismatch base-pair (Y203:OH-dG15:N2 at 3.95 ± 1.58 Å) and its backbone can still interacts with the dG17 phosphate (Y203:N-dG17:P distance of 4.75 ± 1.07 Å). Y203 get also slightly closer to the target strand in 5’ of the damage, exhibiting decreased distances compared to the isolated 8-oxoG: Y203:HH-dT6:O2 at 5.65 ± 2.10 Å and Y203:HH-dA7:O4’ at 5.29 ± 1.82 Å.

A mismatch in 5’ of 8-oxoG induces a slightly different rewiring of the interaction network in the damaged region. N149 interaction with 8-oxoG is also stronger than for an isolated lesion (N149:ND2-8OG:N3 of 3.92 ± 0.94 Å) and hydrogen bonding with the dG17 nucleobase is even more pronounced than with a 3’ mismatch (N149:ND2-dG17:N3 of 3.97 ± 1.18 Å). R154 exhibit similar interaction with the dT19 phosphate than for the 3’ mismatch (5.74 ± 1.22 Å) and get even closer to dG18 backbone (4.67 ± 1.0 Å), but R204 does not get closer to neither dA7 nor dG6 compared to the isolated lesion (R204:CZ-dA7:P at 7.01 ± 2.69 Å and R204:CZ-dG6:N3 at 7.70 ± 1.58 Å). The stable Y203-dG15 interaction observed for the 3’ mismatch is not retrieved with the 5’ mismatch, yet Y203 gets also closer to the dG17 phosphate (Y203:N-dG17:P of 4.62 ± 1.14 Å) and to the dA7 sugar (Y203:HH-dA7:O4’ of 5.13 ± 2.15 Å). The hydrogen bond with the dG6 nucleobase is also tighter than with the isolated lesion (Y203:HH-dG6:O2 at 5.74 ± 2.78 Å).

The stronger interaction networks observed upon the presence of 3’ and 5’ mismatches translates into a stiffening of the protein region harboring the interacting residues. The flexibility profile generated by Machine Learning (ML) analysis of our simulations show higher flexibility values for the reference structure (isolated 8-oxoG) around the N149, R154 and Y203-R204 key amino acids. A global stiffening of the DNA structure is also observed upon addition of a mismatch, regardless of its position. Besides, the clustering analysis reveals a broader distribution of the 4 main clusters in the reference system (25%/22%/20%/15%) than with additional 3’ and 5’ mismatches (44%/20%/12%/8% and 39%/17%/13%/12%, respectively), for which a slightly dominant structure emerges, hence underlying the enhanced stiffness of the hOGG1-DNA - see Figures S1-3 and supplementary pdb files.

#### 3.1.2. Interactions maintaining the DNA-protein interface

At the hOGG1-DNA interface, key-contacts ensure the stability of the complex. Several hOGG1-DNA structures exhibit a hydrogen bonding network involving among others G245, K249 and V250, which are crucial for maintaining the target strand during the 8-oxoG extrusion [14,31–33]. In agreement with these observations, our simulations of the reference system harboring a single 8-oxoG show similar interaction patterns. G245 interacts mostly with dG8 backbone (G245:N-dG8:P distance at 6.22 ± 2.80 Å) and K249 can form hydrogen bonds with dG6 (K249:NZ-dG6:P at 7.71 ± 3.94 Å). V250 does not interact directly with the target strand in our simulations but stays close to the 8-oxoG extrusion path. V250 might interact further in the extrusion process, as we investigate here the ‘encounter’ stage that happens ahead of the extrusion process as described by Shigdel et al. [14]. Interactions with the facing strand are also observed. In line with previous studies, we retrieve contacts between the facing strand in 5’ of the lesion and N151 and R154, mostly with dT19 (N151:ND2-dT19:P at 5.14 ± 1.61 Å and R154:CZ-dT19:P at 6.49 ± 2.33 Å). Our simulations also reveal interactions between K207 and dG8 (K207:NZ-dG8:P at 4.37 ± 1.24 Å), S148 and dA7 phosphate (S148:HG-dA7:P at 4.17 ± 1.87 Å). Interestingly, H270 and K249 lie in the vicinity of 8-oxoG backbone (distance between centers of mass within 12 Å). Hence they remain ideally positioned along the path of the extrusion.

The impact of an additional mismatch on hOGG1 contacts with the DNA backbone appears more pronounced for 5’ than 3’ mismatch position.

The system harboring a 3’ mismatch exhibit the same patterns, yet with a global strengthening of the interactions. The R154 now gets the most interaction with dT19 at the expense of the N151-dT19 contact (N151:ND2-dT19:P at 6.16 ± 2.0 Å and R154:CZ-dT19:P at 5.45 ± 1.46 Å), and K249 and S148 forms more stable hydrogen bonds with the DNA backbone (K249:NZ-dG6:P at 5.60 ± 1.19 Å and S148:HG-dA7:P at 3.13 ± 0.97 Å). On the contrary, the K207-dG8 interaction looses intensity (K207:NZ-dG8:P at 5.17 ± 1.72 Å).

The 5’ position of the mismatch exhibits a drastic stiffening of the hOGG1-DNA backbone interactions. All the contacts already observed in the reference and 3’-mismatched systems get more pronounced, with the G245:N-dG8:P distance dropping at 4.59 ± 1.47 Å, K249:NZ-dG6:P at 5.14 ± 1.34 Å, N151:ND2-dT19:P at 4.63 ± 1.39 Å and R154:CZ-dT19:P at 5.74 ± 1.22 Å, K207:NZ-dG8:P at 3.86 ± 0.56 Å, and S148:HG-dA7:P at 3.61 ± 1.44 Å. Interestingly, G202 and V269 get closer to the DNA backbone upon the presence of clustered lesions, regardless of their position (G202:N-dG17:P at 6.64 ± 1.29 Å and 6.56 ± 1.28 Å, and V269:N-dT6:P at 6.18 ± 1.45 Å and 6.70 ± 1.33 Å for a mismatch in 3’ and 5’, respectively).

### 3.2. DNA structural features

A most important feature for DNA lesion recognition is the structural signature of the damage site within the helix. The perturbation of the structural features of the DNA helix harboring 8-oxoG upon the presence of a vicinal mismatch is described hereafter, focusing on intra- and inter-base pair descriptors and on backbone properties.

#### 3.2.1. Base pair descriptors

The presence of an adjacent mismatch does not strongly perturb the intra-base pair parameters of 8-oxoG - see Figure S4. Yet, a deviation of the base-pair displacement along the X axis can be observed especially with a 5’ mismatch, the average value with the latter being −2.27 ± 0.91 Å while values of −1.38 ± 1.00 Å and −1.76 ± 0.65 Å for the isolated 8-oxoG and with a 3’ mismatch. Likewise, the local inclination of the helix at 8-oxoG slightly increases upon the presence of a mismatch: 5.59 ± 5.0 °, 12.45 ± 4.96 °, 9.45 ± 5.47 ° - see Figure S4. On the contrary, the intra-base pair parameters at the position of the mismatch experience more pronounced deviations, as can be expected from the perturbation of the Watson-crick pairing. Upon a mismatch in 3’ or 5’ to the lesion, the shear parameter at the mismatch base pair increases to around 2.3 Å vs 0.0 Å in the reference - see Figure 5-A. Interestingly, mismatch in 5’ also increases the deviations of the displacement along the X axis by ~ 1 Å at both 3’ and 5’ positions. The local inclinations of the base-pairs in 3’ and 5’of 8-oxoG are slightly impacted by the mismatches, especially when the latter is located in 3’ - see Figures S5 and S6.

Inter-base pairs parameters undergo more drastic perturbations upon the presence of a mismatch, especially between 8-oxoG and its adjacent 5’ base pair - see Figure 5-B. With a mismatch in 5’ of the lesion, the twist angle between the dT(C)4-dG17 / 8-oxoG-dC16 base pairs drops by ~ 10 °. This deviation is of only ~ 3 ° with a 3’ mismatch. A similar trend is observed for the helix twist, denoting a local straightening of the helix due to the 5’ mismatch - see Figure S7. Besides, the distribution of the shift and slide parameters gets much broader with a 5’ mismatch than for the reference system and a 3’ mismatch. Parameters of the 8-oxoG-dC16 / dG(T)6-dC(G)15 base pairs are less impacted by clustered lesions, yet the twist angle increases by ~ 10 ° with a mismatch in 3’ of the lesion. The helix twist angle undergoes a similar deviation with a 3’ mismatch, underlying in this case a more pronounced local deformation of the DNA duplex that might facilitate 8-oxoG extrusion compared to the 5’ mismatch structure - see Figure S8.

#### 3.2.2. Backbone parameters

The structural signature of the 8-oxoG backbone is of utmost importance for its processing by glycosylases [34]. In our simulations, the 8-oxoG ribose puckering is mostly in C1’-exo (50% of the time) and to a lesser extent adopts a C2’-endo (26%) or a O4’-endo (17%) conformation - see Figure 6-A. In presence of an additional mismatch, the 8-oxoG ribose can exhibit a different puckering. When the mismatch is in 3’, the effect is small and the sugar still shows mostly C1’-exo or C2’-endo conformations (40% and 43%). However, with a 5’ mismatch the C3’-exo conformation is much more frequent (24%), yet the puckering is mostly in C1’-exo or C2’-endo (35%and 36%).

The 8-oxoG backbone angles are also different upon clustered lesions, especially with a mismatch in 5’. Distribution of 8-oxoG *ϕ* angle with respect to its *χ* angle show similar trends between the 3’ mismatch and the reference system, but a slightly different pattern when the mismatch is in 5’, the 8-oxoG *χ* angle being then less prone to adopt values around ± 180 ° - see Figure 6-B top. The presence of a 5’ mismatch also strongly impacts the *χ* and *ϕ* angles of the dG6 residue in 3’ of 8-oxoG, with the phase angle (*ϕ*) exhibiting values around 90 ° and *χ* stabilizing around ± 180 °. Indeed, such values are not observed in the reference and the 3’ mismatch systems - see Figure 6-B bottom.

The most pronounced deviations of 8-oxoG backbone values are observed for the *α* and *β* angles. With an isolated 8-oxoG, the *α* angle is mainly centered around −90 ° while *β* lays around 90 °. Upon the presence of a mismatch in 3’, while *α* is still centered around −90 ° and to a lesser extent can adopt values around 90 °, the *β* distribution shows two peaks around 90 ° and ± 180 °. When the mismatch is in 5’ of the lesion, these deviations are more pronounced, with the *α* angle distribution mostly centered around −90 ° and the one for *β* around ± 180 °. The *γ* angle of the 8-oxoG backbone also undergoes non-negligible deviations upon a 5’ mismatch and similar trends are observed for the backbones of the residues in 3’ and 5’ of the lesion - see Figures S9-11.

## 4. Discussion and Conclusions

Understanding how the human DNA repair machinery processes the highly mutagenic 8-oxoG lesion is of crucial importance for cancer research and life sciences. Along with investigations of 8-oxoG formation, structural signature, and processing by the BER enzymes [14,34–38], lower repair yields of 8-oxoG when embedded in local clustered lesions are also a matter of intensive research [6–10]. In this contribution, we explored the molecular mechanisms associated with 8-oxoG recognition by the human hOGG1 glycosylase when 8-oxoG forms clustered lesions with an adjacent mismatch in 5’ or 3’, in order to rationalize their different repair yield, drastically lowered with a 5’ mismatch. Our *μ*s-range simulations allowed us to scrutinize the DNA-protein interaction to delineate perturbations of the key-contacts for the lesion processing, as well as to detail the changes in the structural signature of the DNA duplex which is of utmost importance for its recognition by the repair enzyme.

Investigation of the DNA-protein contacts revealed the perturbation of the interaction network upon the presence of 3’ and 5’ mismatches. In both cases, the lower flexibility of the *α*-helix harboring key residues for the extrusion and of the S148-R154 region harboring the NNN motif conserved in the glycosylases reflect the tighter interactions found upon clustered lesions - see Figure 3. The weakening of the Watson-Crick network might provide more opportunities to form DNA-protein hydrogen bonds upon the presence of a mismatch. Interestingly, with a 5’ mismatch the dG17 is more prone to make interactions with the enzymes and the rewiring of the interactions might perturb the intercalation of Y203 and the interactions involving N149, R204, R154, thus hOGG1 activity. Overall, hOGG1 key-residues are more tightly involved in interactions with the damaged area and with the DNA backbone upon the presence of a mismatch (especially in 5’) than for an isolated 8-oxoG - see Figures 2 and 4. This might be counter-intuitive as DNA duplexes harboring a mismatch or 8-oxoG gets generally more flexible than undamaged oligonucleotides to facilitate the mismatch recognition [39–42]. Hence the importance of exploring the perturbation of their structural behavior when embedded in a clustered lesion, which impacts their repair. The observed global stiffening resulting from the clustered lesions might disfavor the extrusion of the 8-oxoG by hOGG1 by constraining the DNA helix. A process that requires an optimal DNA twisting to efficiently take place as reported for bistranded 8-oxoG / abasic site clustered lesions [43].

**Figure 3.**
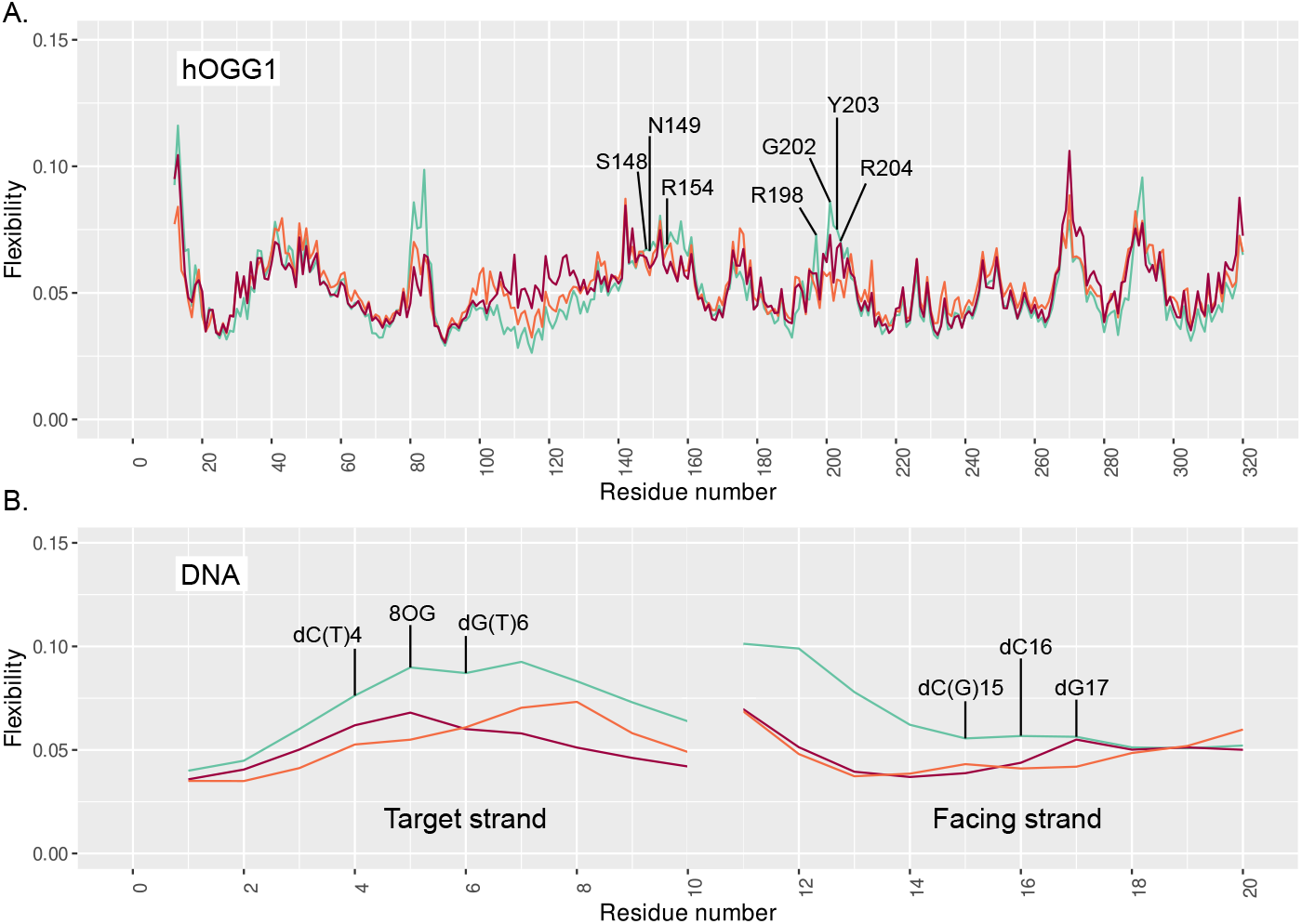
Flexibility profile of hOGG1 (**A.**) and DNA (**B.**) residues, for the reference system in cyan, clustered 8-oxoG + 3’ mismatch in orange and clustered 8-oxoG + 5’ mismatch in red.

**Figure 4.**
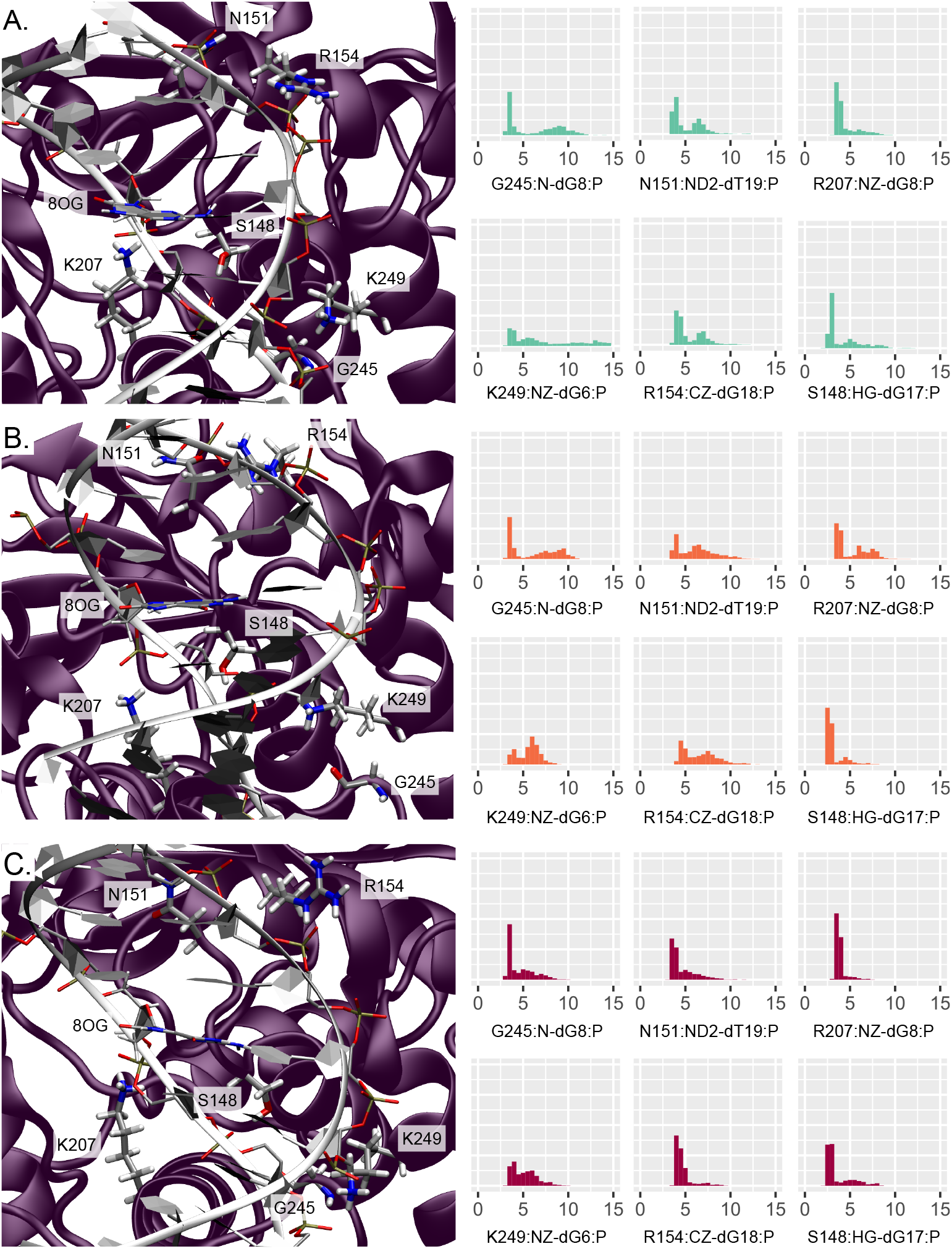
Representation and distribution of the key-interactions with the DNA backbone. **A.** for an isolated 8-oxoG, **B.** for clustered 8-oxoG and 3’ mismatch, **C.** for clustered 8-oxoG and 5’ mismatch. The protein is depicted in purple and the DNA helix in white, the interacting residues side chain are displayed in licorice with thicker tubes for amino acids. The histograms reflect the normalized distribution of the key-distances in Å.

As a matter of fact, our results show that the 5’ mismatch perturbs the geometry of DNA duplex, hence modifying the structural signature of nucleic acid sequence surrounding 8-oxoG - see Figure 5. The impact of a 3’ mismatch is less pronounced, and most of the DNA structural properties exhibit values more comparable to the reference isolated 8-oxoG system. The 8-oxoG backbone angles are particularly deviated by the presence of a 5’ mismatch, and the puckering of the lesion ribose moeity is also perturbed with clustered lesions - see Figure 6. Therefore, the difference of structural signature of the lesion backbone, which is especially important for the recognition and the extrusion by hOGG1, might perturb the lesion processing upon the presence of an adjacent mismatch. Altogether, our simulations delineate the molecular mechanisms driving the difference in repair yields of 8-oxog by hOGG1 when embedded in a clustered lesion with a mismatch: a stiffening of the DNA-protein interactions involving key-residues for the extrusion process, and a modified structural signature of the 8-oxoG and especially in its backbone features, which might disfavor the enzyme processing. Our study brings out a first exploration of the impact of 5’ and 3’ mismatches adjacent to the 8-oxoG on the DNA-protein contacts network and the DNA structural properties. Free energy calculations of the extrusion process would provide complementary information to explore more in-depth the importance of these mechanisms.

**Figure 5.**
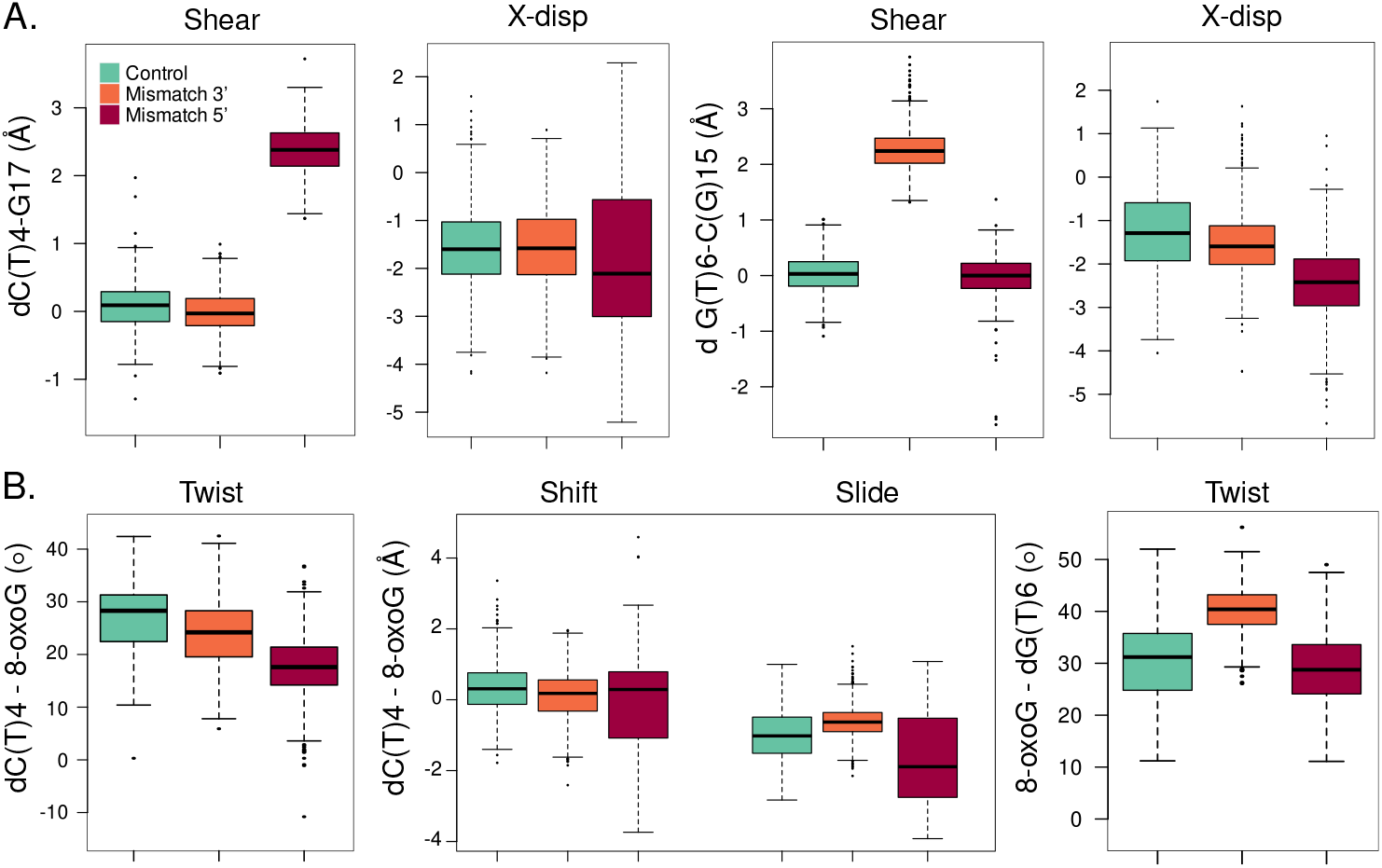
Local structural parameters of the DNA helix harboring an isolated 8-oxoG (cyan), clustered 8-oxoG + 3’ mismatch (orange) or clustered 8-oxoG + 5’ mismatch (red). **A.** Shear and displacement along the X axis (X-disp) intra-base pair parameters for the 5’ dC(T)4-dG17 (left) and 3’ dG(T)6-dC(G)15 base pairs. **B.** Twist, shift and slide inter-base pairs parameters for the dC(T)4-dG17 / 8-oxoG-dC16 base pairs (left) and twist parameter of the 8-oxoG-dC16 / dG(T)6-cC(G)15 base pairs.

**Figure 6.**
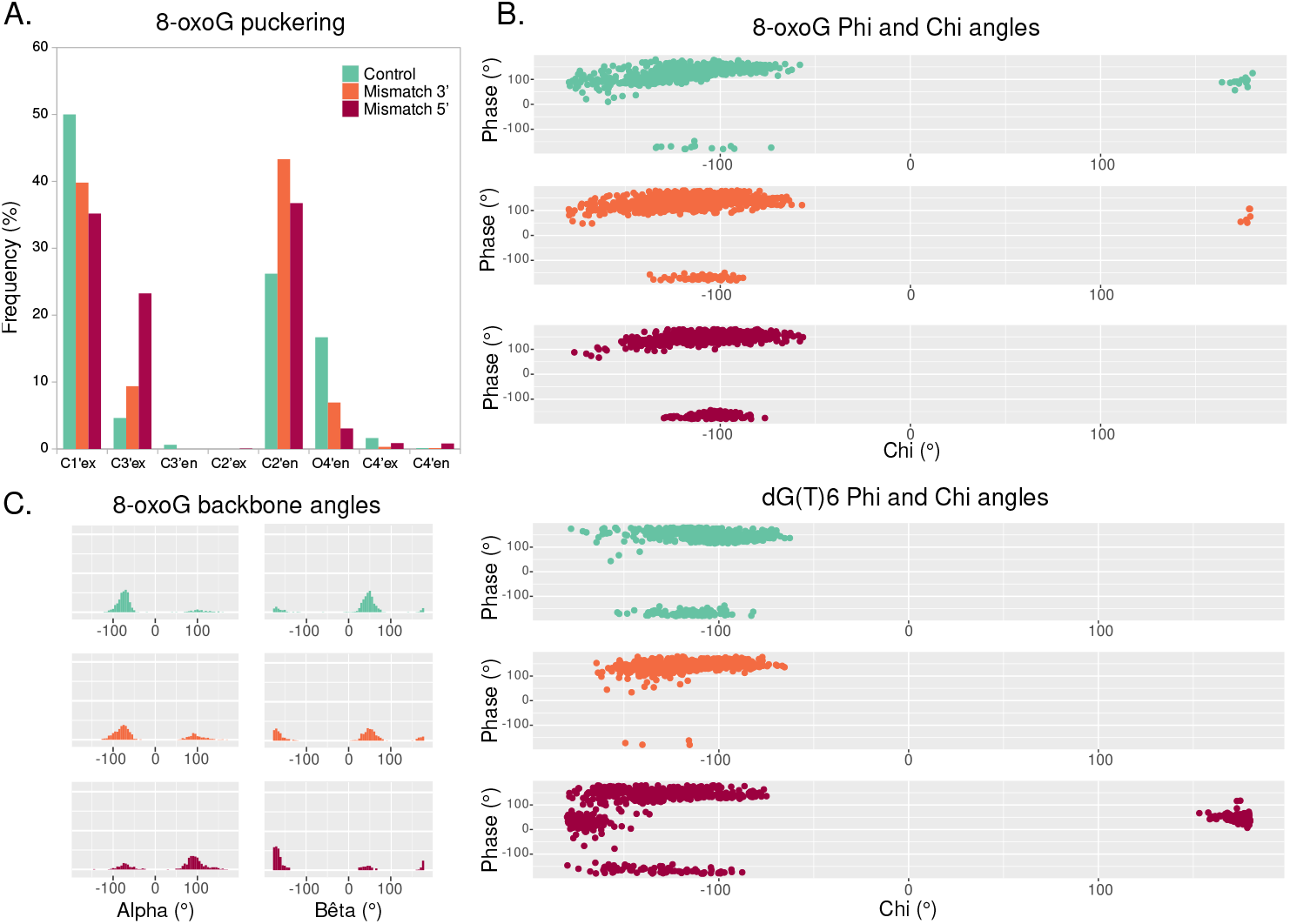
Backbone parameters of the DNA helix harboring an isolated 8-oxoG (cyan), clustered 8-oxoG + 3’ mismatch (orange) or clustered 8-oxoG + 5’ mismatch (red). **A.** Distribution of 8-oxoG sugar puckering. **B.** Phi angle in function of the chi angle of 8-oxoG (top) and dG(T)6 (bottom). **C.** Distribution of 8-oxoG alpha and bêta angles.

## Supplementary Materials

The following are available online, Supp.pdf: supplementary figures, rep-ref-c1-4.pdb : representative structures of the 4 main clusters for the reference system, rep-mm3-c1-4.pdb : representative structures of the 4 main clusters for the system with a mismatch in 3’, rep-mm5-c1-4.pdb : representative structures of the 4 main clusters for the system with a mismatch in 5’, 8OG.targz: AMBER force field parameters used for 8-oxoG.

## Author Contributions

Conceptualization, E.B.; methodology, E.B., T.J., A.M and E.D.; formal analysis, E.B.; data curation, E.B.; writing—original draft preparation, E.B.; writing—review and editing, E.B., A.M and E.D.

## Acknowledgments

The authors thank the regional Explor computing center for computational resources. E.B. thanks the CNRS and French Ministry of Higher Education Research and Innovation (MESRI) for her postdoc fellowship under the “GAVO” program.

## Abbreviations

The following abbreviations are used in this manuscript:

8-oxoG: 8-oxo-7,8-dihydroguanine
hOGG1: human 8-oxoguanine DNA N-glycosylase 1
MD: Molecular Dynamics
ML: Machine Learning

## References

1. ESCODD. Comparative analysis of baseline 8-oxo-7, 8-dihydroguanine in mammalian cell DNA, by different methods in different laboratories: an approach to consensus. Carcinogenesis 2002, 23, 2129–2133.

2. Cadet, J.; Douki, T.; Gasparutto, D.; Ravanat, J.L. Oxidative damage to DNA: formation, measurement and biochemical features. Mutation Research/Fundamental and Molecular Mechanisms of Mutagenesis 2003, 531, 5–23.

3. Gomez-Mendoza, M.; Banyasz, A.; Douki, T.; Markovitsi, D.; Ravanat, J.L. Direct oxidative damage of naked DNA generated upon absorption of UV radiation by nucleobases. The journal of physical chemistry letters 2016, 7, 3945–3948.

4. Maga, G.; Villani, G.; Crespan, E.; Wimmer, U.; Ferrari, E.; Bertocci, B.; Hübscher, U. 8-oxoguanine bypass by human DNA polymerases in the presence of auxiliary proteins. Nature 2007, 447, 606–608.

5. van Loon, B.; Markkanen, E.; Hübscher, U. Oxygen as a friend and enemy: How to combat the mutational potential of 8-oxo-guanine. DNA repair 2010, 9, 604–616.

6. Cadet, J.; Ravanat, J.L.; TavernaPorro, M.; Menoni, H.; Angelov, D. Oxidatively generated complex DNA damage: tandem and clustered lesions. Cancer letters 2012, 327, 5–15.

7. Pearson, C.G.; Shikazono, N.; Thacker, J.; O’Neill, P. Enhanced mutagenic potential of 8-oxo-7, 8-dihydroguanine when present within a clustered DNA damage site. Nucleic acids research 2004, 32, 263–270.

8. Malyarchuk, S.; Youngblood, R.; Landry, A.M.; Quillin, E.; Harrison, L. The mutation frequency of 8-oxo-7, 8-dihydroguanine (8-oxodG) situated in a multiply damaged site: comparison of a single and two closely opposed 8-oxodG in Escherichia coli. DNA repair 2003, 2, 695–705.

9. Malyarchuk, S.; Brame, K.L.; Youngblood, R.; Shi, R.; Harrison, L. Two clustered 8-oxo-7, 8-dihydroguanine (8-oxodG) lesions increase the point mutation frequency of 8-oxodG, but do not result in double strand breaks or deletions in Escherichia coli. Nucleic acids research 2004, 32, 5721–5731.

10. Sage, E.; Shikazono, N. Radiation-induced clustered DNA lesions: Repair and mutagenesis. Free Radical Biology and Medicine 2017, 107, 125 – 135. doi:10.1016/j.freeradbiomed.2016.12.008.

11. Radicella, J.P.; Dherin, C.; Desmaze, C.; Fox, M.S.; Boiteux, S. Cloning and characterization of hOGG1, a human homolog of the OGG1 gene of Saccharomyces cerevisiae. Proceedings of the National Academy of Sciences 1997, 94, 8010–8015.

12. Hill, J.W.; Hazra, T.K.; Izumi, T.; Mitra, S. Stimulation of human 8-oxoguanine-DNA glycosylase by AP-endonuclease: potential coordination of the initial steps in base excision repair. Nucleic acids research 2001, 29, 430–438.

13. Faucher, F.; Doublié, S.; Jia, Z. 8-oxoguanine DNA glycosylases: one lesion, three subfamilies. International journal of molecular sciences 2012, 13, 6711–6729.

14. Shigdel, U.K.; Ovchinnikov, V.; Lee, S.J.; Shih, J.A.; Karplus, M.; Nam, K.; Verdine, G.L. The trajectory of intrahelical lesion recognition and extrusion by the human 8-oxoguanine DNA glycosylase. Nature communications 2020, 11, 1–8.

15. Allgayer, J.; Kitsera, N.; von der Lippen, C.; Epe, B.; Khobta, A. Modulation of base excision repair of 8-oxoguanine by the nucleotide sequence. Nucleic acids research 2013, 41, 8559–8571.

16. David-Cordonnier, M.H.; Boiteux, S.; O’Neill, P. Efficiency of excision of 8-oxo-guanine within DNA clustered damage by XRS5 nuclear extracts and purified human OGG1 protein. Biochemistry 2001, 40, 11811–11818.

17. Kasymov, R.D.; Grin, I.R.; Endutkin, A.V.; Smirnov, S.L.; Ishchenko, A.A.; Saparbaev, M.K.; Zharkov, D.O. Excision of 8-oxoguanine from methylated CpG dinucleotides by human 8-oxoguanine DNA glycosylase. FEBS letters 2013, 587, 3129–3134.

18. Sassa, A.; Beard, W.A.; Prasad, R.; Wilson, S.H. DNA sequence context effects on the glycosylase activity of human 8-oxoguanine DNA glycosylase. Journal of Biological Chemistry 2012, 287, 36702–36710.

19. Phillips, J.C.; Hardy, D.J.; Maia, J.D.; Stone, J.E.; Ribeiro, J.V.; Bernardi, R.C.; Buch, R.; Fiorin, G.; Hénin, J.; Jiang, W.; others. Scalable molecular dynamics on CPU and GPU architectures with NAMD. The Journal of chemical physics 2020, 153, 044130.

20. Case, D.; Ben-Shalom, I.; Brozell, S.; Cerutti, D.; Cheatham III, T.; Cruzeiro, V.; Darden, T.; Duke, R.; Ghoreishi, D.; Gilson, M.; others. AMBER 2018: San Francisco, 2018.

21. Maier, J.A.; Martinez, C.; Kasavajhala, K.; Wickstrom, L.; Hauser, K.E.; Simmerling, C. ff14SB: improving the accuracy of protein side chain and backbone parameters from ff99SB. Journal of chemical theory and computation 2015, 11, 3696–3713. doi:10.1021/acs.jctc.5b00255.

22. Ivani, I.; Dans, P.D.; Noy, A.; Perez, A.; Faustino, I.; Hospital, A.; Walther, J.; Andrio, P.; Goni, R.; Balaceanu, A.; Portella, G.; Battistini, F.; Gelpí, J.L.; González, C.; Vendruscolo, M.; Laughton, C.A.; Harris, S.A.; Case, D.A.;.; Orozco, M. Parmbsc1: a refined force field for DNA simulations. Nature Methods 2016, 38, 55–58. doi:10.1038/nmeth.3658.

23. Bignon, E.; Gillet, N.; Chan, C.H.; Jiang, T.; Monari, A.; Dumont, E. Recognition of a Tandem Lesion by DNA bacterial formamidopyrimidine Glycosylases Explored Combining Molecular Dynamics and Machine Learning. Computational and Structural Biotechnology Journal 2021.

24. Waterhouse, A.; Bertoni, M.; Bienert, S.; Studer, G.; Tauriello, G.; Gumienny, R.; Heer, F.T.; de Beer, T.A.P.; Rempfer, C.; Bordoli, L.; others. SWISS-MODEL: homology modelling of protein structures and complexes. Nucleic acids research 2018, 46, W296–W303.

25. Darden, T.; York, D.; Pedersen, L. Particle mesh Ewald: An N log (N) method for Ewald sums in large systems. The Journal of chemical physics 1993, 98, 10089–10092.

26. Lavery, R.; Moakher, M.; Maddocks, J.H.; Petkeviciute, D.; Zakrzewska, K. Conformational analysis of nucleic acids revisited: Curves+. Nucleic Acids Research 2009, 37, 5917–5929. doi:10.1093/nar/gkp608.

27. Wickham, H. ggplot2: Elegant Graphics for Data Analysis; Springer-Verlag New York, 2009.

28. R Core Team. R: A Language and Environment for Statistical Computing. R Foundation for Statistical Computing, Vienna, Austria, 2017.

29. Humphrey, W.; Dalke, A.; Schulten, K. VMD: visual molecular dynamics. Journal of molecular graphics 1996, 14, 33–38.

30. Fleetwood, O.; Kasimova, M.A.; Westerlund, A.M.; Delemotte, L. Molecular Insights from Conformational Ensembles via Machine Learning. Biophysical Journal 2020, 118, 765–780. doi:https://doi.org/10.1016/j.bpj.2019.12.016.

31. Bruner, S.D.; Norman, D.P.; Verdine, G.L. Structural basis for recognition and repair of the endogenous mutagen 8-oxoguanine in DNA. nature 2000, 403, 859–866.

32. Banerjee, A.; Santos, W.L.; Verdine, G.L. Structure of a DNA glycosylase searching for lesions. Science 2006, 311, 1153–1157.

33. Crenshaw, C.M.; Nam, K.; Oo, K.; Kutchukian, P.S.; Bowman, B.R.; Karplus, M.; Verdine, G.L. Enforced presentation of an extrahelical guanine to the lesion recognition pocket of human 8-oxoguanine glycosylase, hOGG1. Journal of Biological Chemistry 2012, 287, 24916–24928.

34. Qi, Y.; Spong, M.C.; Nam, K.; Banerjee, A.; Jiralerspong, S.; Karplus, M.; Verdine, G.L. Encounter and extrusion of an intrahelical lesion by a DNA repair enzyme. Nature 2009, 462, 762–766. doi:10.1038/nature08561.

35. Lebraud, E.; Pinna, G.; Siberchicot, C.; Depagne, J.; Busso, D.; Fantini, D.; Irbah, L.; Robeska, E.; Kratassiouk, G.; Ravanat, J.L.; others. Chromatin recruitment of OGG1 requires cohesin and mediator and is essential for efficient 8-oxoG removal. Nucleic acids research 2020, 48, 9082–9097.

36. Cadet, J.; Bellon, S.; Berger, M.; Bourdat, A.G.; Douki, T.; Duarte, V.; Frelon, S.; Gasparutto, D.; Muller, E.; Ravanat, J.L.; others. Recent aspects of oxidative DNA damage: guanine lesions, measurement and substrate specificity of DNA repair glycosylases 2002.

37. Dumont, E.; Grüber, R.; Bignon, E.; Morell, C.; Moreau, Y.; Monari, A.; Ravanat, J.L. Probing the reactivity of singlet oxygen with purines. Nucleic acids research 2016, 44, 56–62.

38. Dumont, E.; Grüber, R.; Bignon, E.; Morell, C.; Aranda, J.; Ravanat, J.L.; Tuñón, I. Singlet Oxygen Attack on Guanine: Reactivity and Structural Signature within the B-DNA Helix. Chemistry–A European Journal 2016, 22, 12358–12362.

39. Yang, W. Poor base stacking at DNA lesions may initiate recognition by many repair proteins. DNA repair 2006, 5, 654–666.

40. Sharma, M.; Predeus, A.V.; Mukherjee, S.; Feig, M. DNA bending propensity in the presence of base mismatches: implications for DNA repair. The Journal of Physical Chemistry B 2013, 117, 6194–6205.

41. Buechner, C.N.; Maiti, A.; Drohat, A.C.; Tessmer, I. Lesion search and recognition by thymine DNA glycosylase revealed by single molecule imaging. Nucleic acids research 2015, 43, 2716–2729.

42. Lukina, M.; Koval, V.; Lomzov, A.; Zharkov, D.; Fedorova, O. Global DNA dynamics of 8-oxoguanine repair by human OGG1 revealed by stopped-flow kinetics and molecular dynamics simulation. Molecular BioSystems 2017, 13, 1954–1966.

43. Barone, F.; Dogliotti, E.; Cellai, L.; Giordano, C.; Bjørås, M.; Mazzei, F. Influence of DNA torsional rigidity on excision of 7, 8-dihydro-8-oxo-2-deoxyguanosine in the presence of opposing abasic sites by human OGG1 protein. Nucleic acids research 2003, 31, 1897–1903.

