## Supplementary material for "Molecular mechanisms associated with clustered lesion-induced impairment of 8-oxoG recognition by the human glycosylase OGG1": Supp.pdf

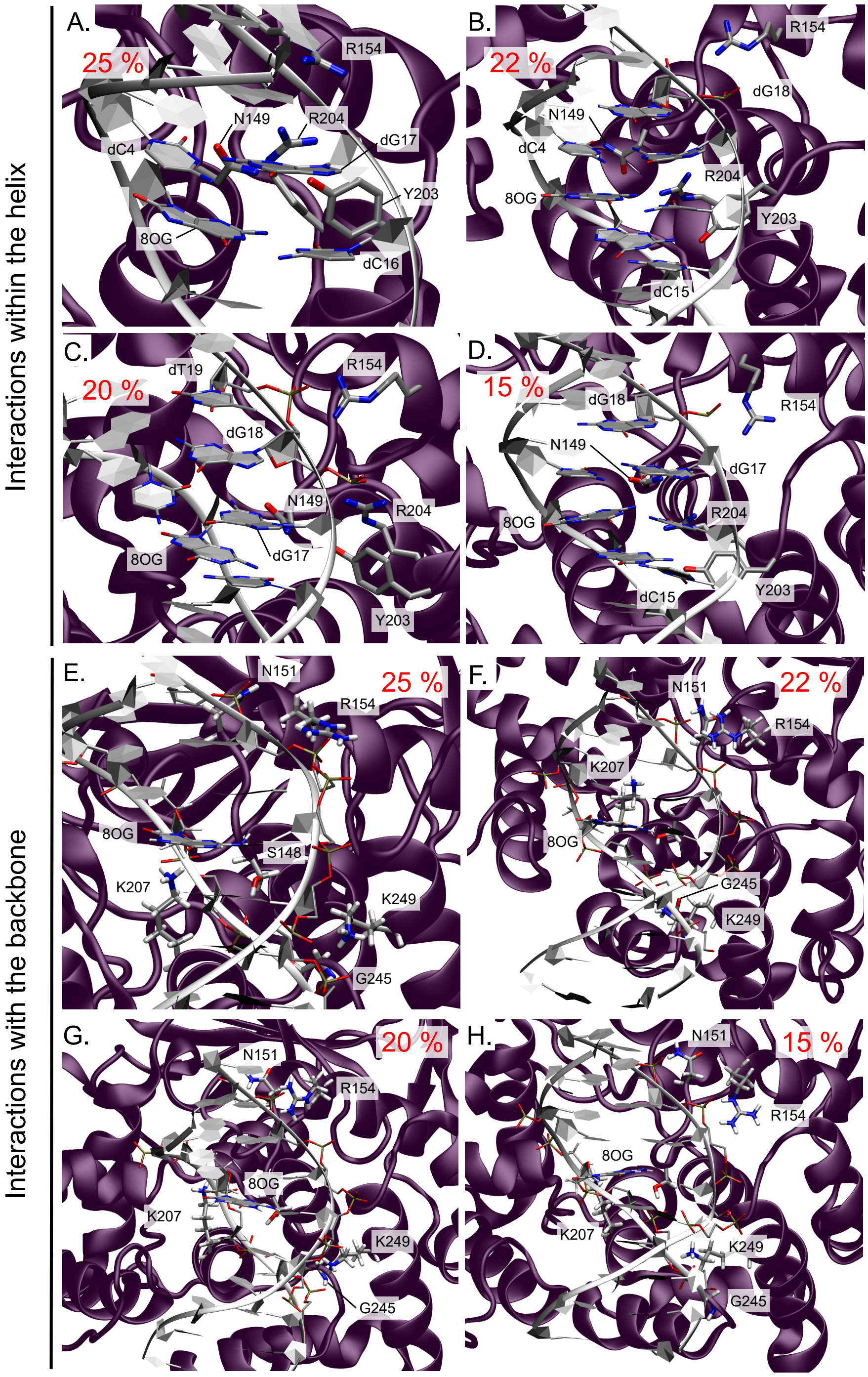


**Figure S1.** Representation of the key-interactions within the DNA helix (**A.** to **D.**) and with the DNA backbone (**E.** to **H.**) upon the presence of an isolated 8-oxoG for the four main clusters. The weight of the cluster is displayed in red.

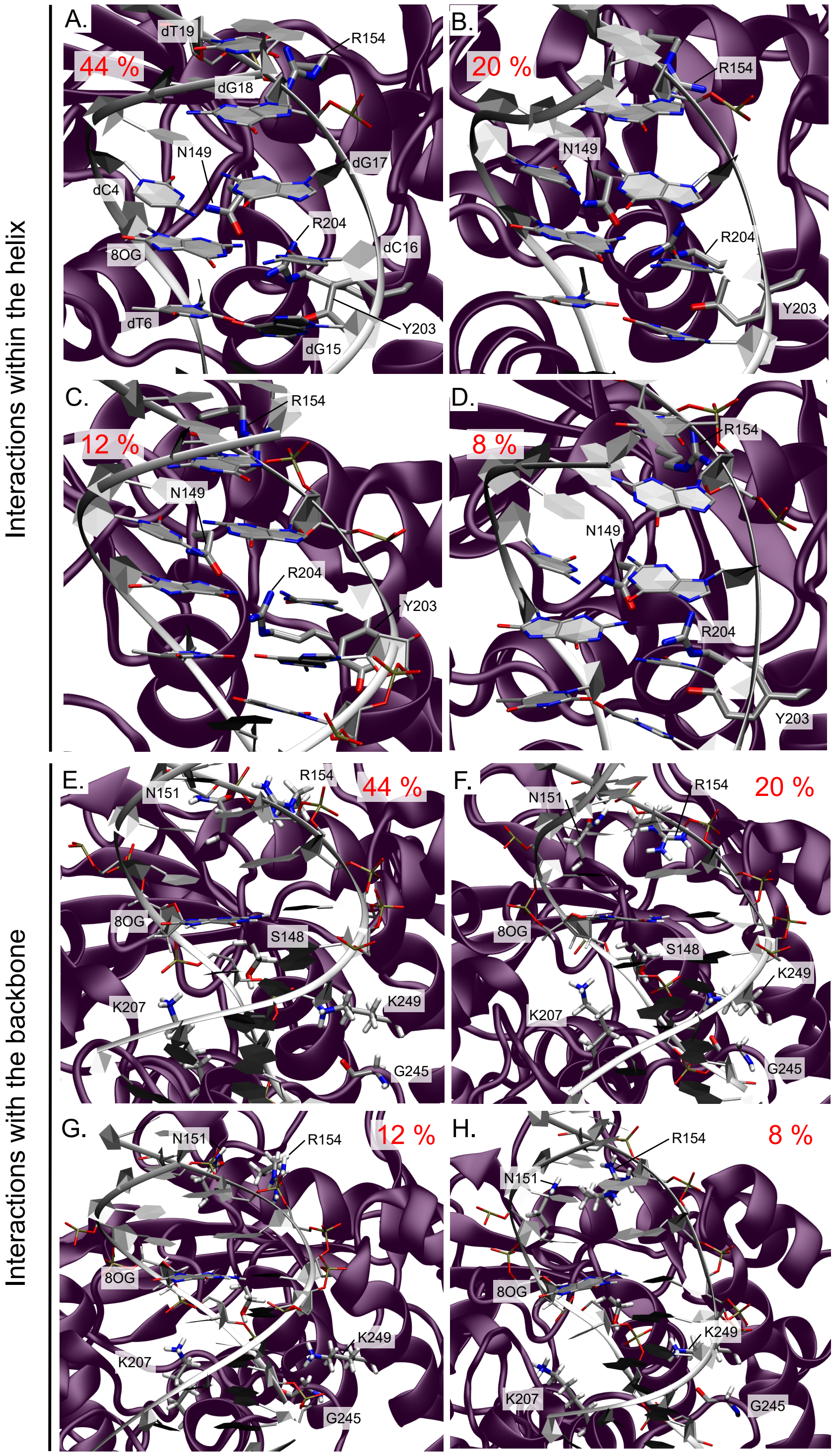


**Figure S2.** Representation of the key-interactions within the DNA helix (**A.** to **D.**) and with the DNA backbone (**E.** to **H.**) upon the presence of clustered 8-oxoG and 3′ mismatch for the four main clusters. The weight of the cluster is displayed in red.

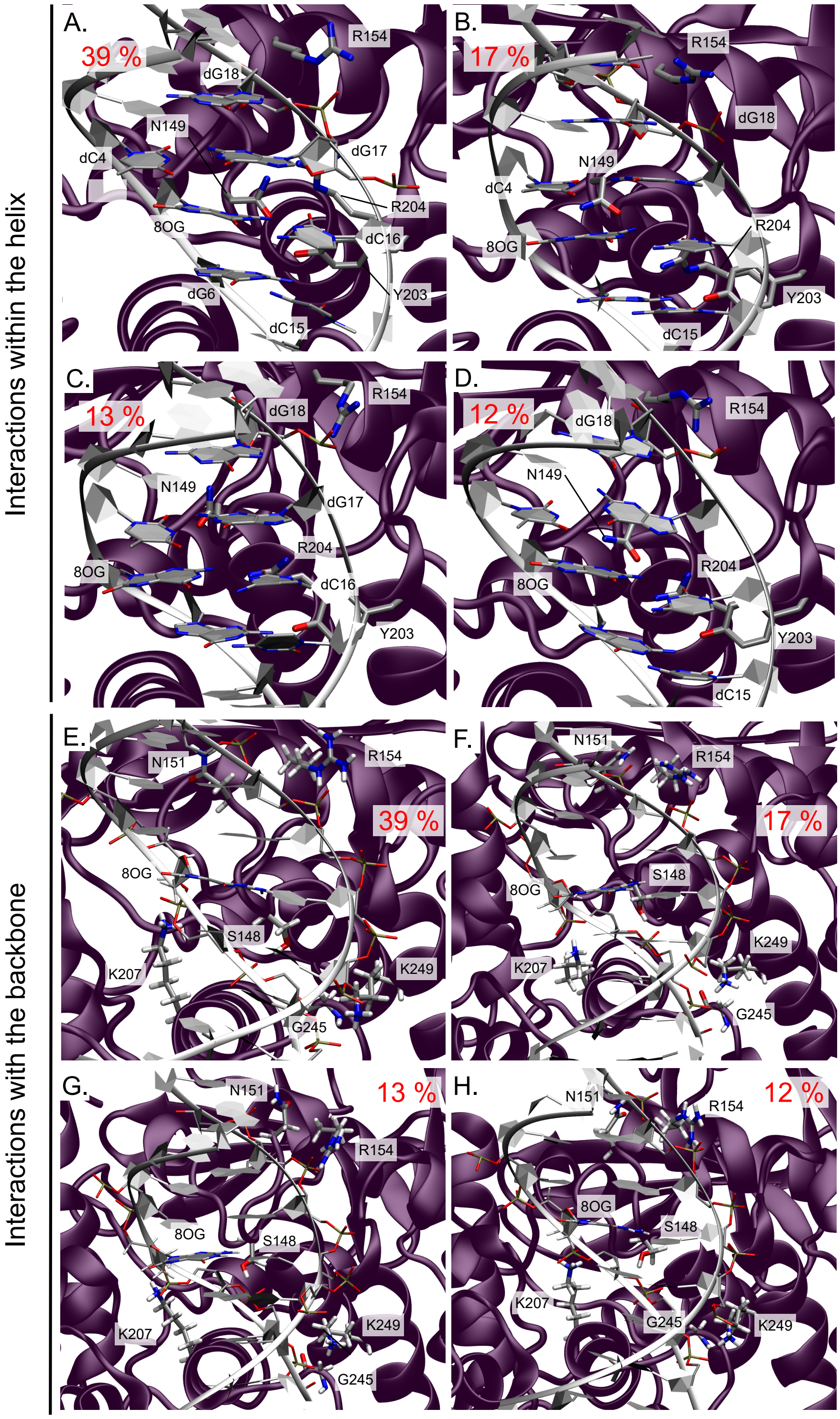


**Figure S3.** Representation of the key-interactions within the DNA helix (**A.** to **D.**) and with the DNA backbone (**E.** to **H.**) upon the presence of clustered 8-oxoG and 5′ mismatch for the four main clusters. The weight of the cluster is displayed in red.

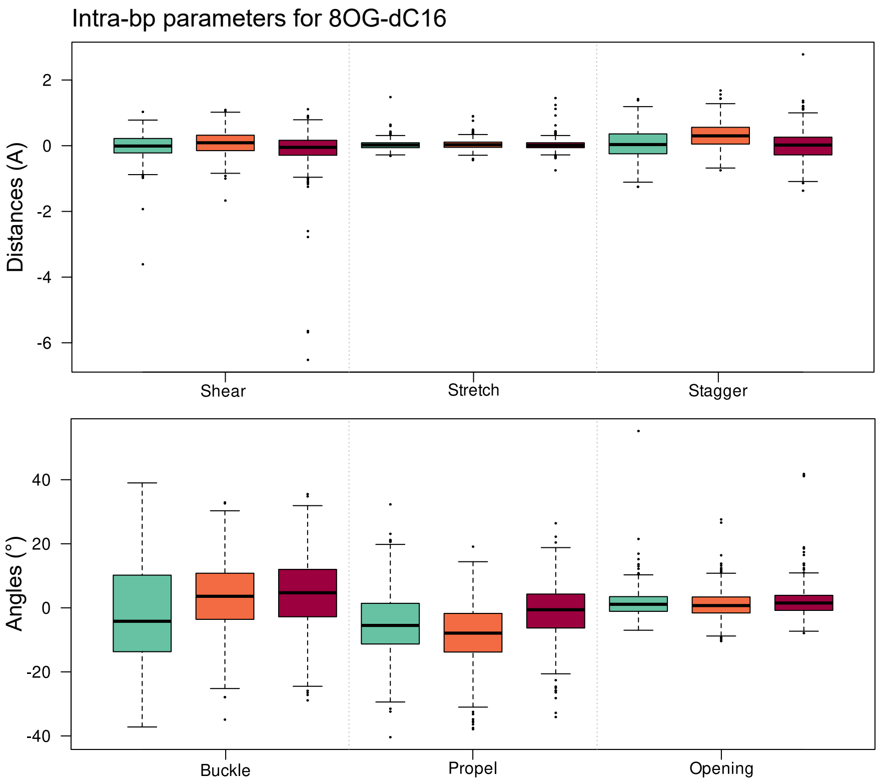


**Figure S4.** Intra-base pair structural parameters of the 8-oxoG-dC16 base pair in systems harboring an isolated 8-oxoG (cyan), clustered 8-oxoG + 3′ mismatch (orange) or clustered 8-oxoG + 5′ mismatch (red).

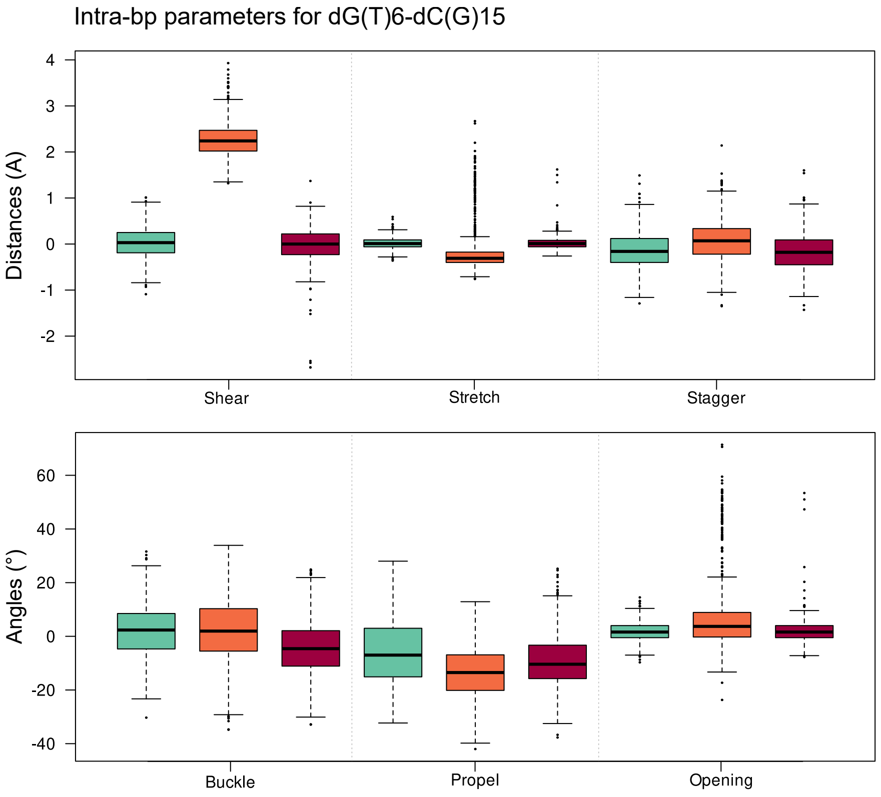


**Figure S5.** Intra-base pair structural parameters of the dG(T)6-dC(G)15 base pair 3′ to the 8-oxoG in systems harboring an isolated 8-oxoG (cyan), clustered 8-oxoG + 3′ mismatch (orange) or clustered 8-oxoG + 5′ mismatch (red).

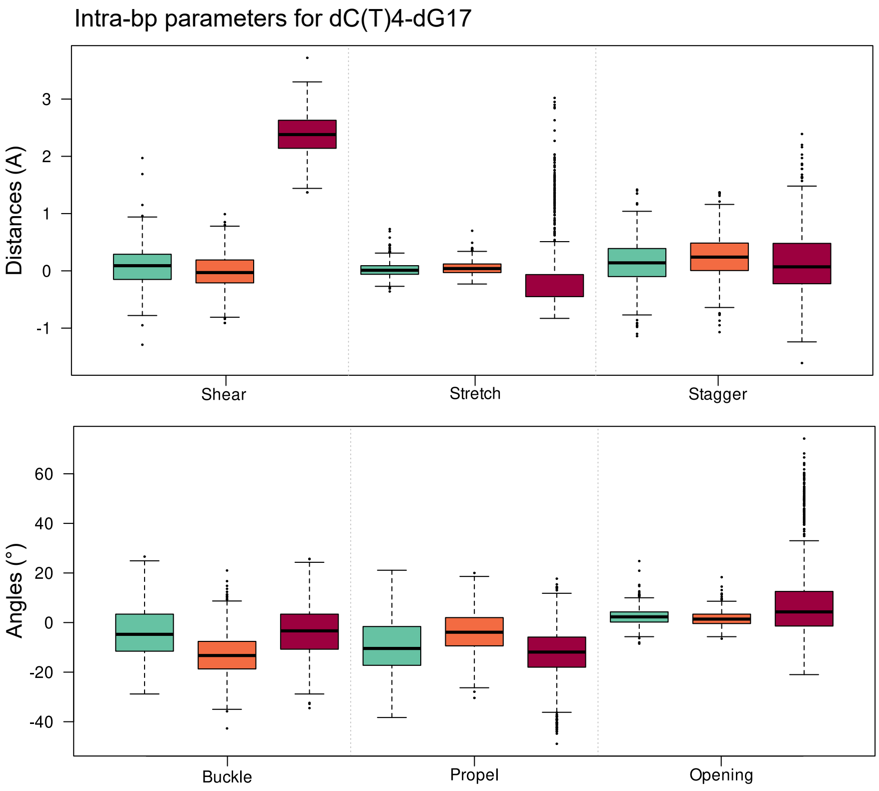


**Figure S6.** Intra-base pair structural parameters of the dC(T)4-dG17 base pair 3′ to the 8-oxoG in systems harboring an isolated 8-oxoG (cyan), clustered 8-oxoG + 3′ mismatch (orange) or clustered 8-oxoG + 5′ mismatch (red).

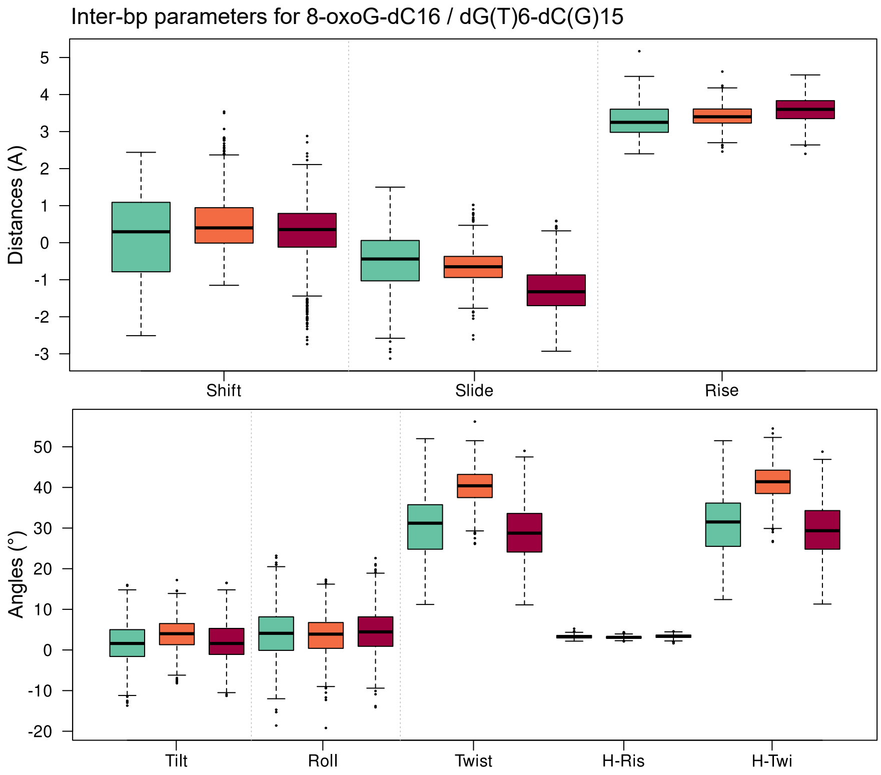


**Figure S7.** Inter-base pairs structural parameters of the 8-oxoG-dC16 / dG(T)6-dC(G)15 base pairs in systems harboring an isolated 8-oxoG (cyan), clustered 8-oxoG + 3′ mismatch (orange) or clustered 8-oxoG + 5′ mismatch (red).

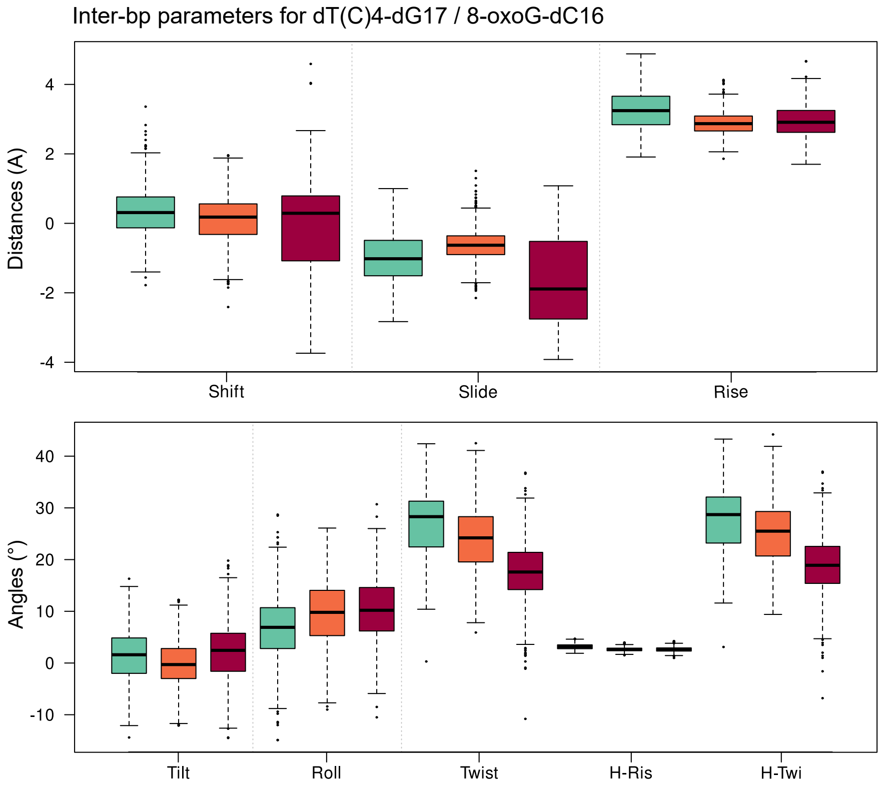


**Figure S8.** Inter-base pairs structural parameters of the dC(T)4-dG17 / 8-oxoG-dC16 base pairs in systems harboring an isolated 8-oxoG (cyan), clustered 8-oxoG + 3′ mismatch (orange) or clustered 8-oxoG + 5′ mismatch (red).

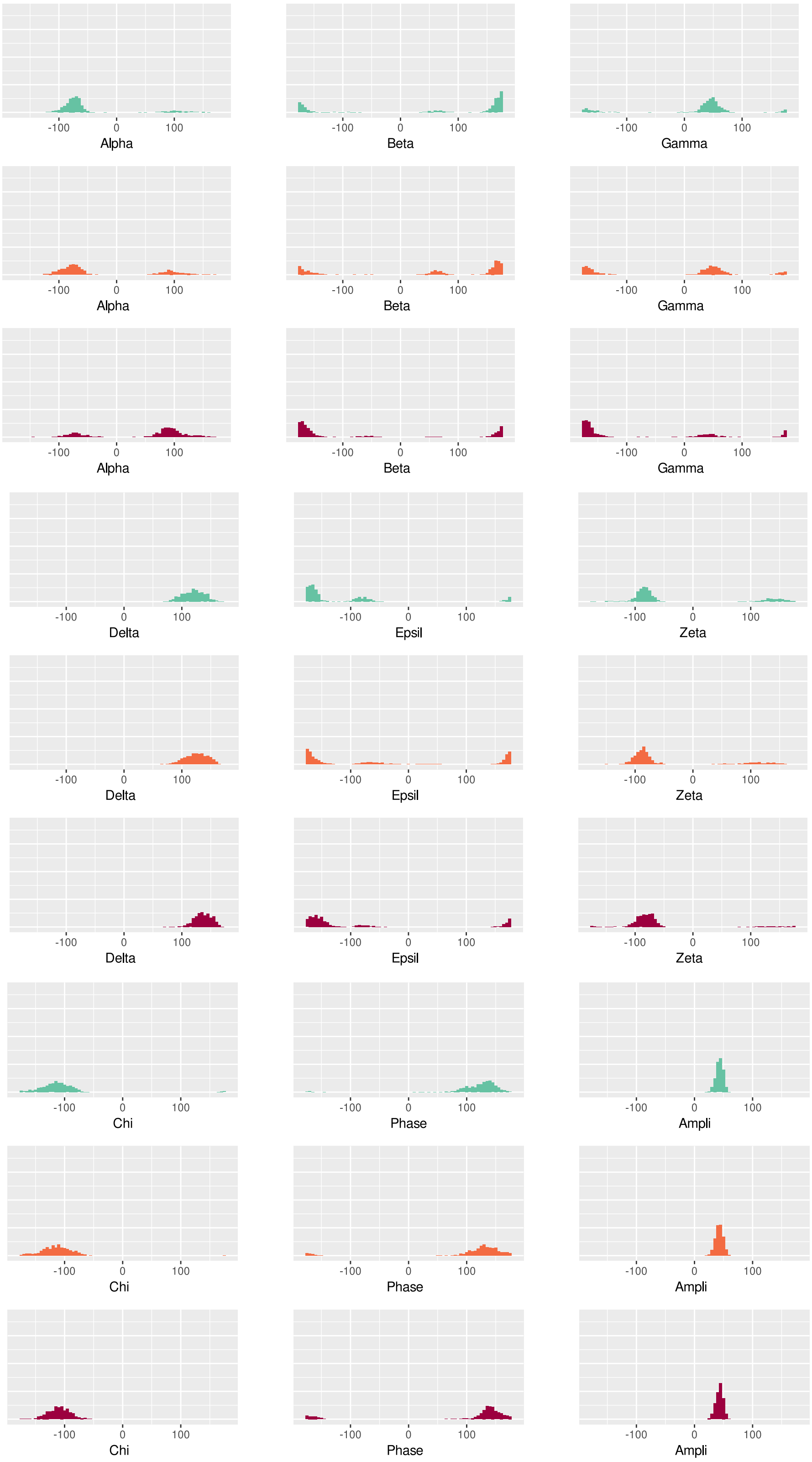


**Figure S9.** Distribution of 8-oxoG backbone angles values in systems harboring an isolated 8-oxoG (cyan), clustered 8-oxoG + 3′ mismatch (orange) or clustered 8-oxoG + 5′ mismatch (red).

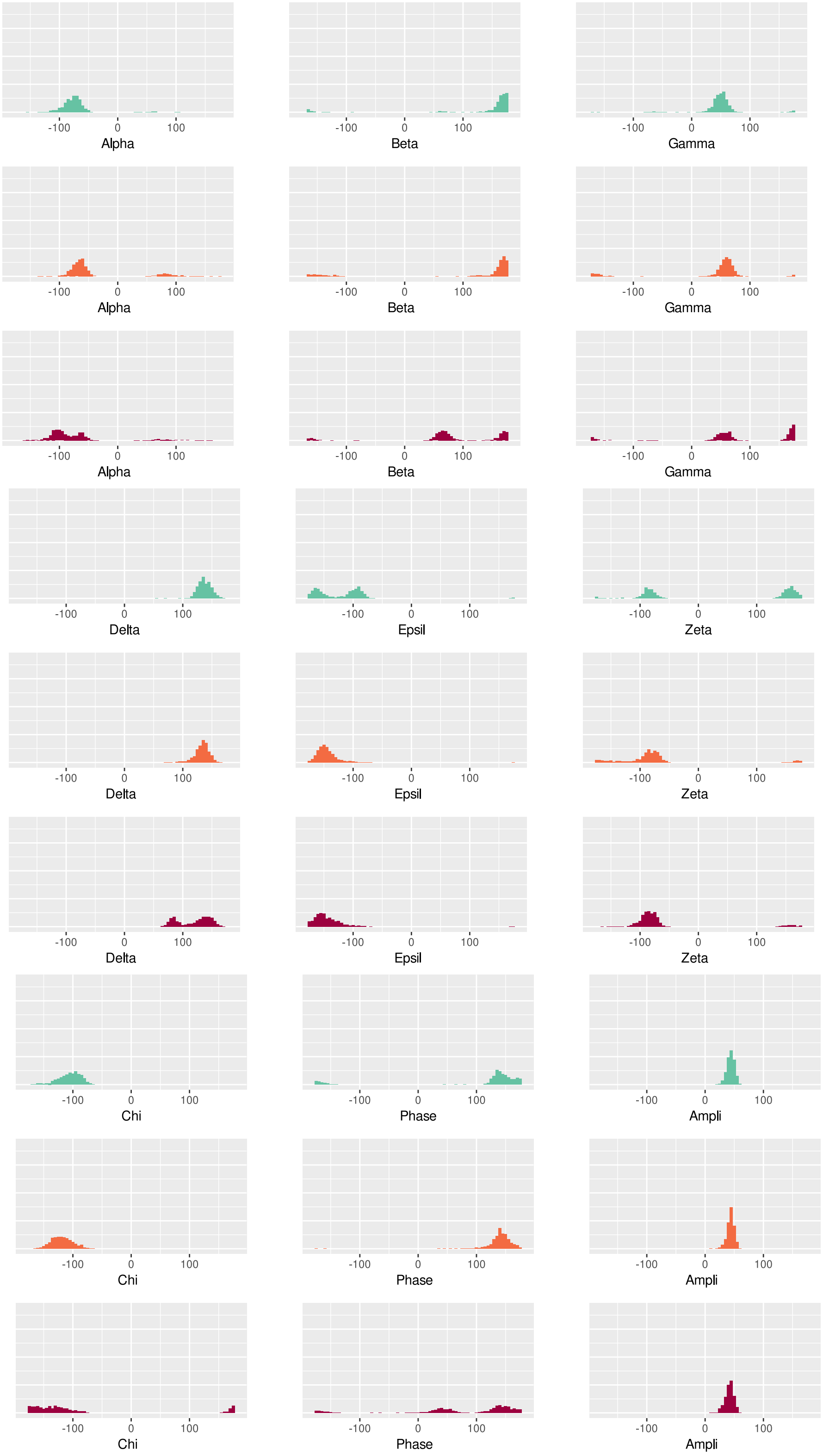


**Figure S10.** Distribution of dG(T)6 backbone angles values in systems harboring an isolated 8-oxoG (cyan), clustered 8-oxoG + 3′ mismatch (orange) or clustered 8-oxoG + 5′ mismatch (red).

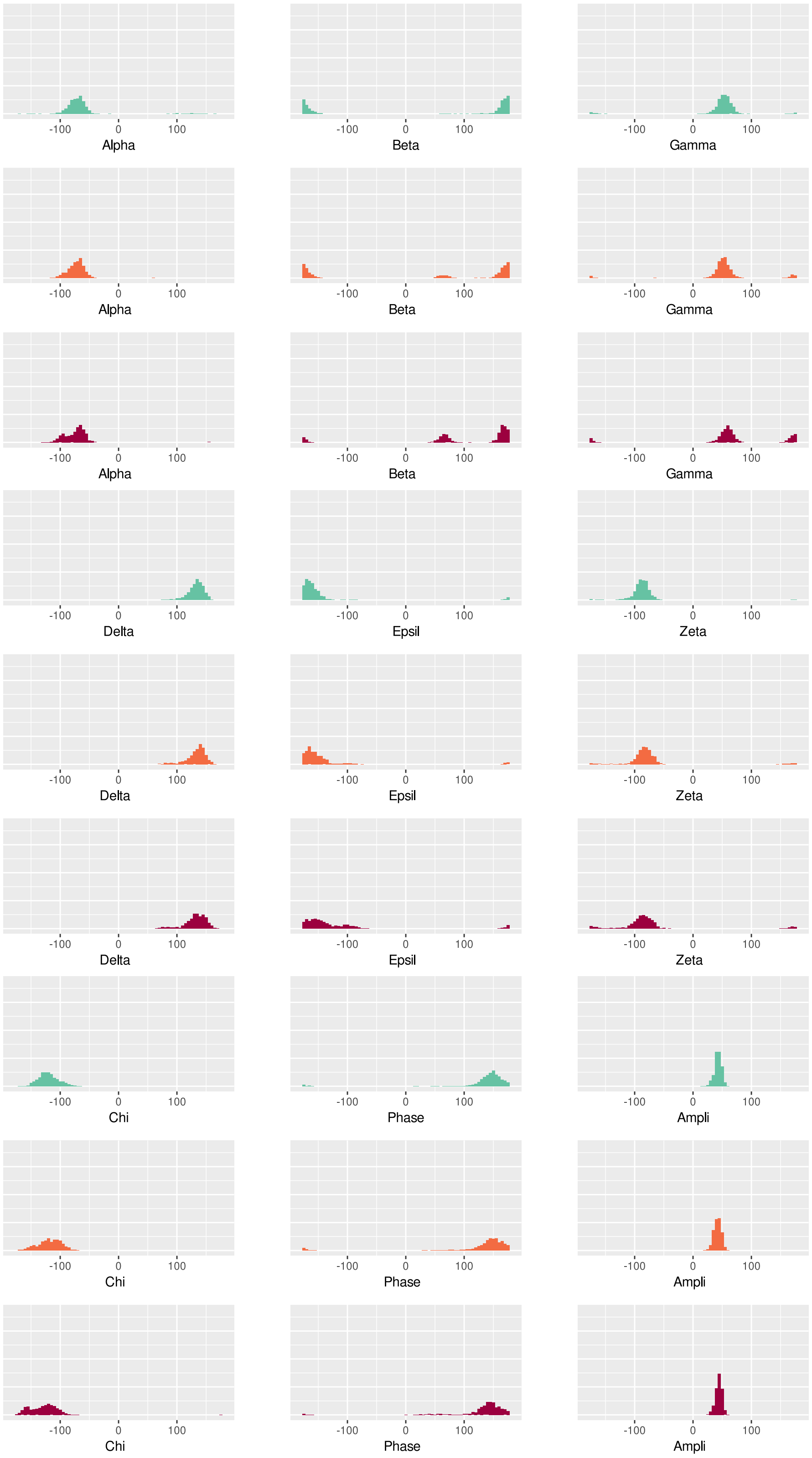


**Figure S11.** Distribution of the dC(T)4 backbone angles values in systems harboring an isolated 8-oxoG (cyan), clustered 8-oxoG + 3′ mismatch (orange) or clustered 8-oxoG + 5′ mismatch (red).
